# Probing DNA - transcription factor interactions using single-molecule fluorescence detection in nanofluidic devices

**DOI:** 10.1101/2021.05.12.443786

**Authors:** Mattia Fontana, Šarūnė Ivanovaitė, Simon Lindhoud, Willy van den Berg, Dolf Weijers, Johannes Hohlbein

## Abstract

Single-molecule fluorescence detection offers powerful ways to study biomolecules and their complex interactions. Here, we combine nanofluidic devices and camera-based, single-molecule Förster resonance energy transfer (smFRET) detection to study the interactions between plant transcription factors of the auxin response family (ARF) and DNA oligonucleotides that contain target DNA response elements. In particular, we show that the binding of the unlabelled ARF DNA binding domain (ARF-DBD) to donor and acceptor labelled DNA oligonucleotides can be detected by changes in the FRET efficiency and changes in the diffusion coefficient of the DNA. In addition, our data on fluorescently labelled ARF-DBDs suggest that, at nanomolar concentrations, ARF-DBDs are exclusively present as monomers. In general, the fluidic framework of freely diffusing molecules minimizes potential surface-induced artifacts, enables high-throughput measurements and proved to be instrumental in shedding more light on the interactions between ARF-DBDs monomers and between ARF-DBDs and their DNA response element.

## Introduction

Single-molecule techniques are playing an increasingly important role in the investigation of properties and dynamics of proteins and biomolecular complexes thanks to their ability to resolve inter- and intramolecular heterogeneity. In particular, fluorescence-based techniques became widely employed thanks to their ease of use (Joo et al., 2008). Traditionally, single-molecule fluorescence detection (SMFD) can be carried out either on a confocal microscope, which uses one or more avalanche photodiodes as point detectors, or on a wide-field microscope used in total-internal reflection fluorescence (TIRF) mode, which uses emCCD or sCMOS cameras to monitor hundreds of molecules in parallel. SMFD of free-diffusing molecules on a confocal microscope allows for high time resolution (typically *µ*s) at the expense of throughput and short observation times while SMFD of surface-immobilized molecules on a TIRF microscope displays a somehow complementary behaviour with lower time resolution (typically ms (Farooq and Hohlbein, 2015)) compensated by high throughput and long observation times.

During the past decade, different frameworks were proposed to overcome the limitations imposed by these traditional implementations of SMFD. For confocal microscopy, the main focus has been prolonging the observation times (Kim et al., 2011; Wunderlich et al., 2013; Tyagi et al., 2014; Kim et al., 2015) while in TIRF-based applications, the aim was to eliminate the need of sample immobilization (Leslie et al., 2010; Fontana et al., 2019; Gilboa et al., 2019). Performing SMFD experiments on a TIRF microscope without immobilization allows to minimize surface-induced artifacts whilst maintaining the high throughput inherent to camera-based detection schemes.

Here, we utilize a glass-made fluidic device (Fontana et al., 2019) on a TIRF microscope to study some members of the transcription factor family of auxin response factors (ARFs) and their interaction with their DNA response element. The ARF family of transcription factors is involved in the response to auxin, a plant hormone that regulates many developmental processes. While much is known about their binding preferences, a detailed understanding of the binding process is still lacking. We were able to detect the interaction between ARF DNA binding domain (ARF-DBD) with a doubly labelled DNA construct containing the response element by monitoring the change in its FRET signature and the diffusion coefficient upon DNA binding. We then studied fluorescently labelled ARF-DBDs to show that they are present in their monomeric form at the nanomolar concentrations used in our experiments, which contrast with the dimeric form portrayed in multiple crystal structures (Boer et al., 2016; Kato et al., 2020).

## Materials and Methods

### Device fabrication, surface passivation and cleaning

The fluidic devices were fabricated by Micronit (Micronit Microtechnologies B.V., The Netherlands) following the same methodology and design described previously (Fontana et al., 2019). To prevent non-specific adsorption of the analytes, we opted for passivating cleaned nanochannels using polyethylene glycol (mPEG, MW = 5000 Da, Laysan Bio Inc., USA) as follows. First, the channels were washed with acetone and then incubated for 5 minutes with a 50:1 acetone:Vectabond solution; then, channels were washed with acetone (1X) followed up by MilliQ (3X) and incubated with a solution of mPEG in MOPS buffer for 3 hours. The pegylated channels were then washed with PBS buffer and stored in a wet chamber at 4 °C. After the experiments, the devices were washed by flowing and incubating a solution of 50:1 MilliQ:Hellmanex®III (Hellma, Germany). After rinsing with MilliQ, the devices were burned in a furnace at 500 °C.

### Protein expression and purification

The DBDs of MpARF1, MpARF2, AtARF1 and AtARF5 were amplified and cloned in a modified pTWIN1 vector. The expression and purification of recombinant ARF-DBDs were performed as previously described (Boer et al., 2016; Kato et al., 2020).

### Protein labelling

The labelling buffer is composed of 20 parts of solution 1 (274 mM NaCl, 5.4 mM KCl, 10 mM phosphate) and one part of solution 2 (0.2 M NaHCO_3_, pH adjusted to 9 using NaOH); the pH of the final labelling buffer was adjusted to 8.3 using solution 1 or 2. Buffer exchange was carried out by two consecutive runs on Zeba™Spin Desalting Columns 7MWCO (Thermo Fisher Scientific, USA) that was equilibrated in labelling buffer. For labelling ARF5-DBD, 60 µL of a 3.5 mg mL^−1^ ARF5-DBD solution was supplemented with 12 µL of 800 µM DNA duplex and 1 mM TCEP (final protein concentration: 65 µM, final DNA concentration: 133 µM). 5 µL of 10 mM label was added. The labelling reaction was carried out overnight at 4 °C; then, the labeled protein was separated from free dye on a PD-10 desalting column (Cytiva, USA) that was equilibrated in PBS supplemented with 5 mM DTT. For ARF1-DBD, the labelling was performed as described for ARF5-DBD but in absence of the DNA duplex.

### Accessible Volume simulations

The web server 3D-DART (van Dijk and Bonvin, 2009) was used to first model a straight, standard B-DNA structure for the oligo used in the single-molecule FRET (smFRET) experiments. A second, bent oligo was modelled using the geometrical information extracted from the short DNA present in the crystal structure of MpARF2 (Kato et al., 2020). The bent oligo was then rigid-docked in place of the short oligo that was present in the crystal structure. Both oligos were then used as starting points to model the accessible volumes of the FRET pair using the FRET-restrained positioning and screening software (Kalinin et al., 2012). The accessible volume is defined by the region of space that each dye can explore given its own geometry, the geometry of the linker and the attachment point on the biomolecule (see table 1).

**Table 1.** AV simulations: Geometrical parameters of the dyes taken from reference Craggs et al. (2019).

| Dye | Linker | L. length<br>[Å] | L. width<br>[Å] | R<sub>1</sub><br>[Å] | R<sub>2</sub><br>[Å] | R<sub>3</sub><br>[Å] |
| --- | --- | --- | --- | --- | --- | --- |
| Cy3B | C6-NHS | 14.2 | 4.5 | 8.2 | 3.3 | 2.2 |
| ATTO647N | C6-NHS | 17.8 | 4.5 | 7.4 | 4.8 | 2.6 |

### Single-molecule detection and tracking

Single-molecule detection was performed using a TIRF microscope (previously described (Fontana et al., 2019)) equipped with a fiber-coupled laser engine (Omicron, Germany). The triggering of the lasers and the camera as well as the laser intensities were controlled by a home-written LabVIEW program. The power of the lasers was set to 100 % which results in a measured laser power at the fiber output of approximately 280 mW for the green excitation (561 nm) and 140 mW for the red excitation (638 nm). The illumination was based on the stroboscopic alternating-laser excitation (sALEX (Farooq and Hohlbein, 2015)) scheme with laser pulses of 1.5 ms in a frame time of 10 ms. Green and red pulses of two subsequent frames were placed back-to-back to facilitate tracking between these coupled frames. Particles were localized using a modified version of GaussStorm (Holden et al., 2010) and tracked using Maria Kilfoil’s MATLAB porting of a tracking algorithm developed in IDL (Crocker and Grier, 1996). For samples containing only one fluorophore (e.g. singly-labelled proteins), a scheme based on stroboscopic back-to-back illumination was applied to the single excitation laser. Per condition, we acquired 4 movies of 4000 frames for a total acquisition time of 160 s.

### FRET and ALEX

The apparent FRET efficiency *E*\* was calculated from the emission intensities of donor and acceptor after donor excitation (denoted as *DD* and *DA*) according to

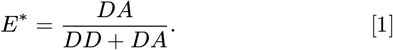

Moreover, alternating laser excitation (ALEX) scheme was used; in this scheme every frame where the donor is excited is followed by one of direct excitation of the acceptor fluorophore using a second laser resulting in a third photon stream (*AA*) for each molecule (Kapanidis et al., 2004; Hohlbein et al., 2014; Hellenkamp et al., 2018). This additional information allows for calculating the stoichiometry ratio *S*, defined as

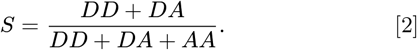

*S* can be used to filter molecules: molecules with a stoichiometry close to 0 have no photoactive donor whereas molecules with a stoichiometry close to 1 have no photoactive acceptor. A stoichiometry around 0.5 represents molecules having both photoactive donor and acceptor.

### Particle displacements analysis

The particle displacements between two back-to-back illuminated frames were obtained using 2D Gaussian fitting of the imaged point spread functions as described previously (Fontana et al., 2019). From the variance of the 1D Gaussian distributions of all one-dimensional displacements in either *x*, or *y*-direction (*σ*^2^, also called mean-square-displacement *MSD*), we calculated the mean particle diffusion coefficient *D* using

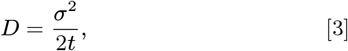

here, *t* is the time between the mean time of two back-to-back illuminations (1.5 ms). In case the displacements are calculated along the direction of the flow, the mean flow speed (⟨*v*⟩) is determined by the mean (*µ*) of their distribution

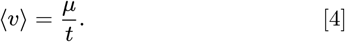

We note that equations 3 and 4 do not take into consideration the uncertainty associated with the localization. While equation 4 holds true, calculating the diffusion coefficient from 3 requires knowledge on the variance that was generated by the localization uncertainty (of both localizations) and its subtraction from the total variance to isolate the component that was generated by the Brownian motion. Consequently, diffusion coefficients calculated without accounting for the localization uncertainty will overestimate the real value.

### Calculation of theoretical diffusion coefficients

The predicted diffusion coefficients presented in this chapter were calculated using HYDROPRO (Ortega et al., 2011). This software uses the crystal structure of a macromolecule to estimate its diffusion coefficient. The calculations were performed loading the pdb file of the protein or complex of interest, setting the temperature to 20 °C and the viscosity of the solution to 1 cP; the calculations were performed in *Mode 1* (shell-mode from atomic-level) using a radius of the atomic elements of 2.84 Å.

## Results

### SMFD in parallel nanochannels

Detecting non-immobilised, nanometer-sized molecules in fluidic devices using camera-based microscopy is challenging as the movement of the molecule due to diffusion and advection within the time of a single camera frame results in the spreading of emitted photons over many pixels. Given the already limited photon budget of single emitters, the movement further decreases the signal-to-noise ratio (SNR).

To mitigate the motion blur caused by advection, the fluidic design should provide sufficiently slow flow speeds while still allowing for fast exchange of sample solutions ensuring low dead times. A convenient way of implementing such characteristics is to use parallel flow control (PFC (Liang et al., 2008; Mathwig et al., 2012)). In our implementation of PFC (Fontana et al., 2019), the flow coming from a syringe pump is divided between the imaging channels (i.e. nanochannels) and a bypassing microchannel (Fig. 1a-b) such that most of the liquid passes through the microchannel due to its lower hydraulic resistance thereby reducing the fluid velocity inside the nanochannels by several orders of magnitude (to values in the tens or hundreds of nanometres per millisecond). This arrangement further assures that the dead volume in the tubing and feeding channel is replaced in minutes (i.e. reduced dead time). The nanochannels have an height of 200 nm thereby limiting the movement of the fluorescent molecule to the evanescent field of the TIRF microscope and enabling reliable single-particle tracking (Fig. 1b). Another source of spreading of the emitted photons over additional pixels is the movement of the fluorescent molecule due to diffusion; the effect can be strongly reduced by applying a stroboscopic illumination scheme in which the molecule is illuminated only for a fraction of the acquisition time of the frame (Farooq and Hohlbein, 2015). Moreover, the illumination of two neighbouring frames can be arranged in a back-to-back configuration (Fig. 1c) to further minimize the displacements between those frames thus allowing for higher concentrations of molecule in the channels before successful tracking is compromised by overlapping pathways.

**Fig. 1.**
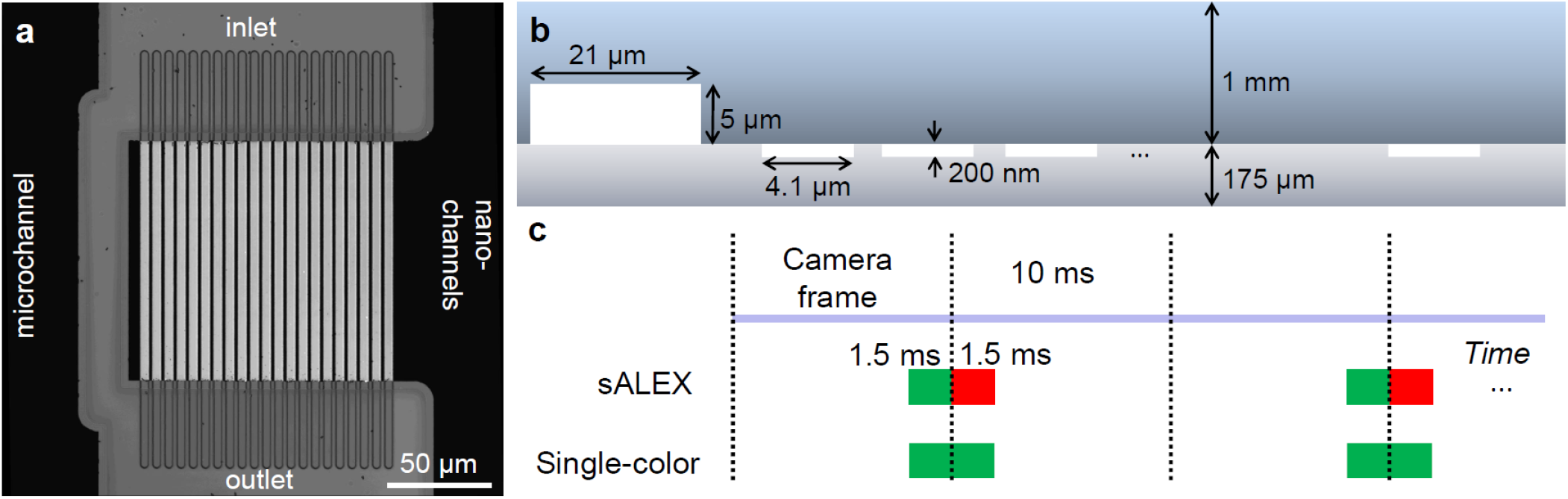
Design of the fluidic device and excitation (Adapted from reference Fontana et al. (2019)). (a) High-resolution confocal reflection scan of the parallel nanochannels device; both the array of imaging nanochannels (l × w × h: 120 by 4.1 by 0.2 µm) and the bypassing microchannel (l x w × h: 120 by 21 by 5 µm) are visible. (b) Schematic cross-section of the parallel channel array, showing the dimensions of the microchannel and the nanochannels. The microchannel has a larger cross-section thereby causing lower resistance to the fluid. As a result, most of the sample liquid passes through the microchannel minimising both the speed of the analyte inside the nanochannels and the dead time of the fluidic system. (c) Schematic representation of the laser excitation schemes. The concept of stroboscopic illumination is used to reduce the motion blur of single-particle localization while the excitation of two successive frames are placed back-to-back to facilitate tracking and allowing for higher concentrations of fluorescent molecules to be analysed.

### The interaction between ARF and labelled DNA can be visualized using smFRET in nanochannels

We tested the feasibility of visualizing the interaction between ARF and its DNA response element in nanofluidic devices by flowing a DNA construct through the channels in presence or absence of ARF transcription factors. We utilized a doubly-labelled DNA construct similar to the one we previously used as immobilized entity in single-molecule FRET experiments to determine the binding affinity of the DNA Binding Domains (DBDs) of *Marchantia polymorpha* ARF 1 and 2 (Kato et al., 2020). The construct contains two high affinity AuxREs (TGTCGG) in an inverted topology with a spacer of 7 bp (IR7) and is labelled with a FRET pair featuring Cy3B as the donor and ATTO647N as the acceptor fluorophore (Fig. 2a). The accessible volumes of the dyes, as well as the average distances between each other, were calculated using the FPS software (Kalinin et al., 2012) (see also Materials and Methods). Upon binding to DNA, ARF is expected to bend the DNA and sterically confine the movement of the dyes (Fig. 2b); thus, ARF binding causes a decrease in the FRET efficiency of the DNA construct (Kato et al., 2020). Measuring the DNA construct alone (2.5 nM in PBS buffer containing oxygen scavenger system (Rasnik et al., 2006; Cordes et al., 2009): 1 mM Trolox, 1% gloxy and 1% glucose) in the fluidic devices let to a single population centered at *E*\* = 0.39 (Fig. 2c, top); a second sample containing saturating concentration (256 nM) of MpARF2-DBD showed the characteristic shift toward a lower FRET efficiency (*E*\* = 0.31, Fig 2c, bottom).

**Fig. 2.**
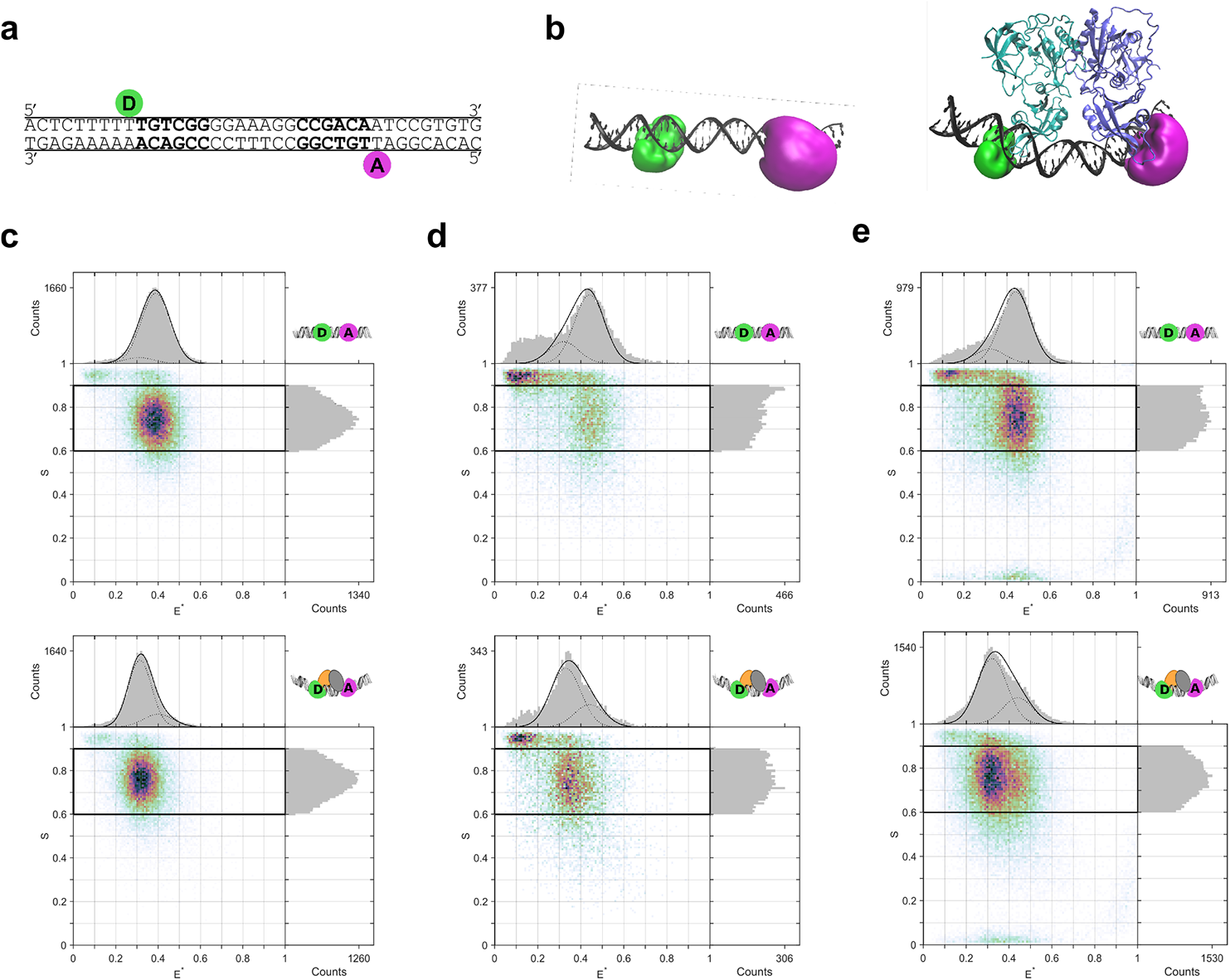
Visualizing ARF-DNA interactions in nanofluidic devices. (a) Schematic representation of the doubly-labelled DNA construct with the two AuxREs (bold) and the positions of the donor (D, Cy3B) and the acceptor (A, ATTO647N) fluorophore. (b) Accessible Volume (AV) simulation for free (left) and ARF-bound (right) DNA construct. The AV represents the volume that can be visited by the dye thanks to its flexible linker. In presence of ARFs, the mean distance between the dyes increases resulting in a decrease of the FRET efficiency *E*\*. (c) *E*\**S* histograms of a sample containing DNA alone (top) and DNA with a saturating concentration of MpARF2-DBD (256 nM, bottom). The fraction of bound DNA can be calculated from the relative areas of the two Gaussians centered at the *E*\* of the free and bound DNA. (d) The same samples of (c) were loaded consecutively on the syringe while being separated by a air bubble, and without the standard enzymatic oxygen scavenger system being present. The shift in *E*\* upon ARF binding can be seen, but the reduced data quality due to photobleaching does not allow to reliably identify the relative amount of bound and free DNA. (e) Same experiment as in (d) but with MpARF1-DBD instead of MpARF2-DBD.

We asked to which extend the use of parallel nanochannels allows performing titration experiments without the supervision of an operator. To this end, we opted for sequentially loading the samples containing DNA and increasing concentration of ARF inside the syringe or, alternatively, inside a long connecting tubing. To prevent mixing of neighbouring samples, we separated them by an air bubble. This arrangement abolishes dispersing effects created by convection (i.e. Taylor dispersion) while the air bubbles provide a visual cue for the position of the sample. We tested the general feasibility by loading two samples separated by an air bubble into a syringe before feeding the nanochannels. The first sample contained the DNA construct in PBS 1X buffer (2.5 nM, Fig. 2d,e top) while the second contained both DNA and 256 nM of either MpARF2-DBD (Fig. 2d, bottom) or MpARF1-DBD (Fig. 2e, bottom). The ARF containing samples showed the characteristic shift towards lower FRET efficiencies when compared to the one without ARF; however, the noise present on the *E*\**S* histogram makes the evaluation of the DNA bound fraction inaccurate.

The time duration of an entire titration combined with the presence of air bubbles between the samples makes the use of enzymatic oxygen scavenging system (OSS) problematic; given the presence of molecular oxygen in the sample, we explain the lower quality of the data to be caused by a relevant portion of the counts in the *E*\**S* histogram originating from doubly-labelled DNA molecules in which either the donor or the acceptor fluorophore photobleached during the acquisition of a single frame.

Taken together, these results show that the characteristic shift in FRET efficiency seen in experiments with the surface-immobilised DNA construct can be also seen using freely diffusing DNA. This finding further confirms that the shift in *E*\* is caused by specific ARF to DNA binding rather than surface-induced artifacts (e.g. protein absorption on the surface).

### Changes in the diffusion coefficient of labelled DNA can be used to monitor ARF-DNA interaction

In the previous section we described experiments in which doubly-labelled single DNA constructs were imaged and tracked to determined their FRET efficiency (*E*\*) in absence or in presence of ARF; the resulting change in *E*\* was then used as readout for ARF binding. In this section, we will analyse the change in diffusion coefficient (*D*) of the DNA constructs to obtain an independent second readout for binding.

During two back-to-back illuminations, the DNA construct in the nanochannels will move due to flow and diffusion. In the direction perpendicular to the flow, the movement is due exclusively to Brownian motion and the distribution of the displacements can be used to calculate the diffusion coefficient (*D* = *σ*^2^*/*2*t*, see also Materials and methods). The diffusion coefficient of the DNA construct in 1x PBS buffer (without OSS) was found to be 62 and 67 µm^2^ s^−1^ in two independent experiments (Fig. 3a,b, top) and thereby close to the theoretical one of 69.8 µm^2^ s^−1^ (see Materials and Methods). Upon addition of either MpARF1-DBD or MpARF2-DBD, the diffusion coefficient of the DNA construct decreases to 52 µm^2^ s^−1^ and 54 µm^2^ s^−1^ respectively (Fig. 3a,b, bottom); these values are again consistent with the theoretical value for the diffusion coefficient of the complex formed by a dimer of MpARF2-DBD bound to the DNA construct which is 51.0 µm^2^ s^−1^. The analysis of the diffusion coefficients in presence of an OSS shows decreased diffusion coefficients for both free DNA and bound DNA. This effect is likely to be caused by (I) the reduced photobleaching which in turn reduces the average localization error and (II) the increased viscosity of the sample when OSS are present in the solution. (e.g. 1% glucose = +2% viscosity (Haynes, 2016)). Similar to the experiments without the OSS, addition of ARF decreased the diffusion coefficient of the DNA by around 12 µm^2^ s^−1^.

**Fig. 3.**
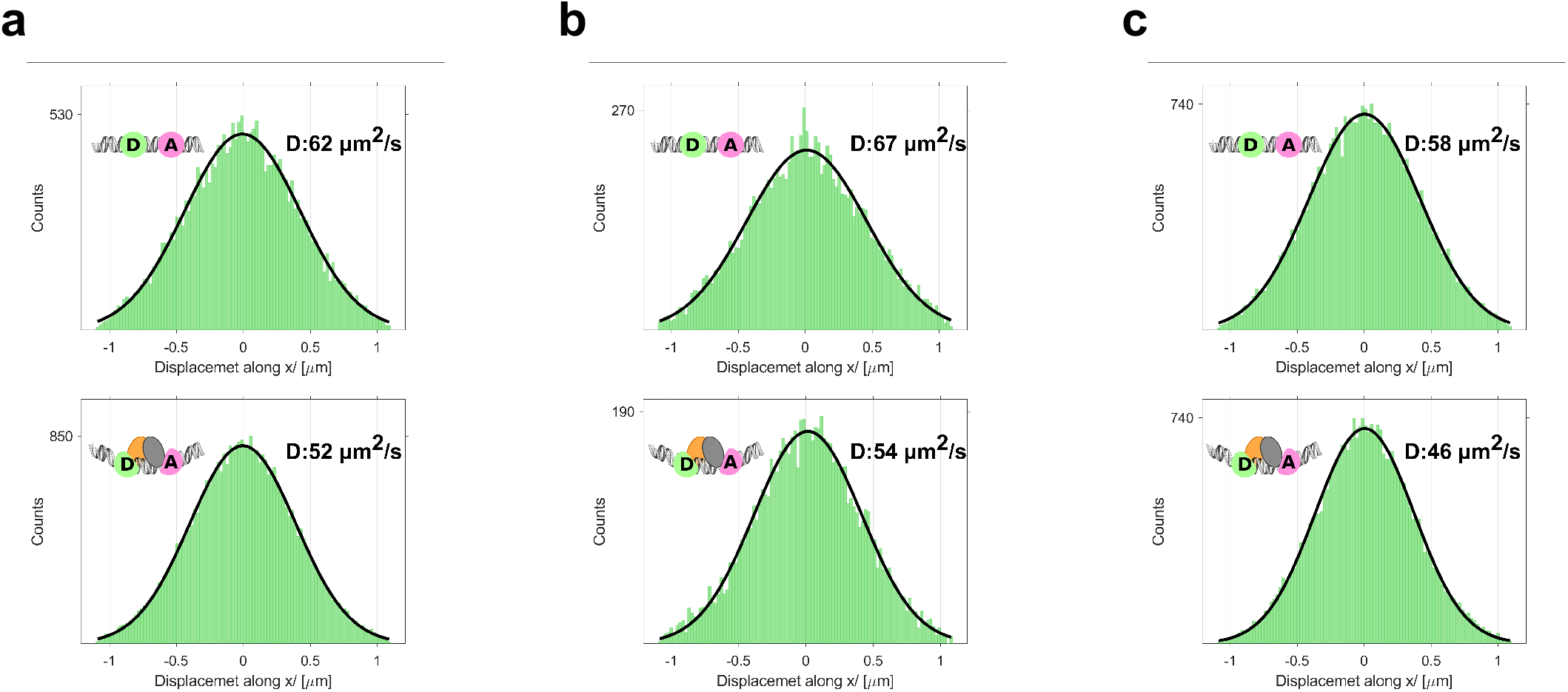
Displacement analysis of the labelled DNA construct in absence or presence of MpARF1-DBD and MpARF2-DBD. (a) The diffusion coefficient of the DNA construct decreases by 10 µm^2^ s^−1^ when 256 nM of MpARF1-DBD is added to the solution, consistent with the formation of a complex of the DNA construct with a protein dimer. (b) Same experiment as (a) but using MpARF2-DBD instead of MpARF1-DBD; the starting and final diffusion coefficients are in good agreement with the results in (a). (c) Same experiment as in (b) but with the addition of an enzymatic oxygen scavenger system (Glucose Oxidase/Catalase, glucose and Trolox); the decrease in diffusion coefficient upon MpARF2-DBD addition is consistent with (a) and (b) but the absolute values are lower due to the decrease in localization error and an increase of the viscosity of the solution.

### ARF-DBDs at nanomolar concentration are monomeric

A common feature of ARF-DBDs as portrayed in crystal structures is that they form dimers caused by interactions between their dimerization domains (DD). Unfortunately, the structures do not provide information on the dimerization equilibrium in solution. To the best of our knowledge, the only data available (acquired using SAXS, Small Angle X-ray Scattering) reported ≈ 60% dimer when 78 µM *A. thaliana* ARF1-DBD was added in solution (Boer et al., 2016). This places a dissociation constant of the dimer to be ∼ 10 µM. Given the scarcity of information about ARF dimerization in solution we decided to perform experiments in parallel nanochannels. We imaged stochastically labelled AtARF1-DBD and AtARF5-DBD (≈ 1 nM) whilst applying the back-to-back stroboscopic illumination scheme (Fontana et al., 2019); the proteins were tracked and the distributions of displacements were determined (Fig. 4).

**Fig. 4.**
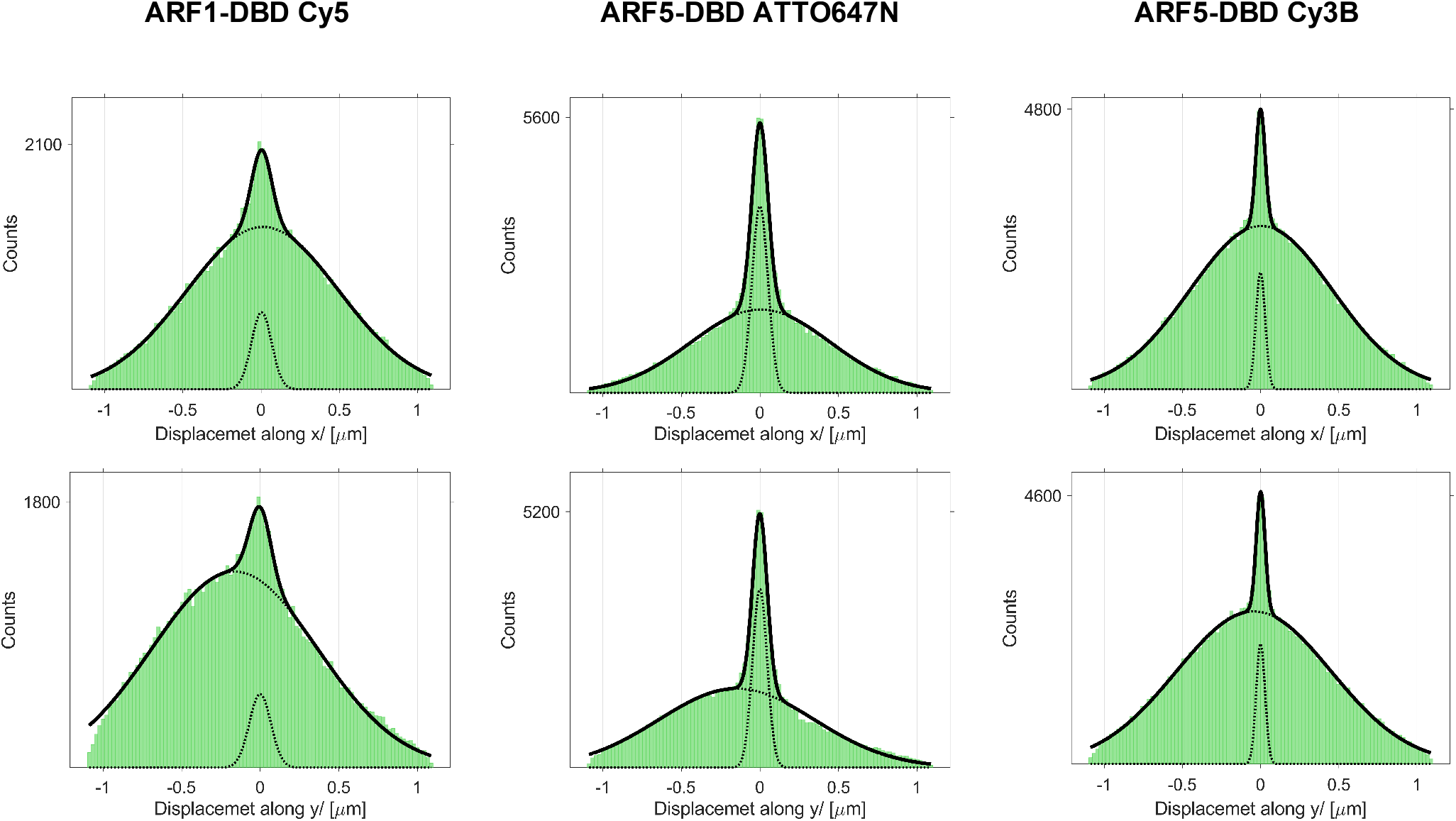
Displacements analysis for AtARF1-DBD (Cy5), AtARF5-DBD (ATTO647N) and AtARF5-DBD (Cy3B). The displacements perpendicular to the flow (along *x*, top) and parallel to the flow (along *y*, bottom) are best fitted with two Gaussians. The slow population (narrow Gaussian) belongs to the proteins absorbed to the surface of the channel as it appears centred around zero even along the direction of flow. The width of the Gaussian fit for this population equals double the localization error and was used to correct the diffusion coefficient *D* of the freely moving population (wide Gaussian). The resulting diffusion coefficient along *x* was 78 µm^2^ s^−1^ for AtARF1-DBD labelled with Cy5, 65 µm^2^ s^−1^ for AtARF5-DBD labelled with ATTO647N and 69 µm^2^ s^−1^ for AtARF5-DBD labelled with Cy3B.

Despite the passivisation of the glass surfaces with PEG, a small but noticeable fraction (*<* 20%) of single-molecule localization came from ARFs that were adsorbed on the surface; for this reason a two Gaussian fit was employed, in which one Gaussian fits the displacements of the free flowing fraction and the second one the apparent displacements of the adsorbed population. Fitting the distribution of displacements perpendicularly to the direction of flow (along *x*, Fig. 4, top) resulted in two Gaussian distributions centred at 0. The wider Gaussian belongs to the free-flowing molecules and its width is proportional to the diffusion coefficient of the free-flowing species while the narrow one belongs to the adsorbed population and its width is equal to double the localization error for an immobile particle (see Materials and methods). An apparent diffusion coefficient for the adsorbed population was calculated and subtracted from the one of the free-moving particles to account for the localization error; we note that although moving particles are expected to have a higher localization error due to motion blur, we used stroboscopic illumination which minimizes this contribution. We obtained diffusion coefficients of 78 µm^2^ s^−1^ for AtARF1-DBD labelled with Cy5, 65 µm^2^ s^−1^ for AtARF5-DBD labelled with ATTO647N and 69 µm^2^ s^−1^ for AtARF5-DBD labelled with Cy3B. These values can be compared with the theoretical ones for monomeric and dimeric AtARF5-DBD (extracted from the crystal structures PDBID:4LDU, see Materials and methods) of 73 µm^2^ s^−1^ and 55 µm^2^ s^−1^ respectively (given the high similarity in structure and size, the values obtained for AtARF1-DBD using the crystal structure PDBID:4lDV showed no difference when rounded to two significant digits). The comparison of the simulated and experimental values for the diffusion coefficient suggests that, at the concentration tested (≈ 1 nM), both AtARF1-DBD and AtARF5-DBD are present as monomers in solutions. Fitting the distribution of displacements along the direction of the flow (along *y*, Fig. 4, bottom) proved that the fraction with a low apparent diffusion coefficient corresponded to absorbed molecules and not proteins aggregates. Here the narrow distribution is still centred around zero while the wide population of the freely-moving particles is centred around a value which is proportional to the mean flow speed in the field of view (⟨*v*⟩, see Materials and methods). In general, the lack of a narrower population centred around the mean flow speed proves that protein aggregation, if present, is negligible. We note that the diffusion coefficient of the moving particle along *y* is affected by additional broadening due to the inhomogeneous velocity field inside the channels and cannot be used to determine the diffusion coefficient directly.

## Conclusion

In this work, we used single-molecule fluorescence detection performed in fluidic devices to study the DNA binding domain of several members of the family of auxin response factor (ARF) transcription factors and the interaction between them and their DNA response element.

We found that the interaction between doubly-labelled ds-DNA bearing the response element and unlabelled ARF led to changes in both the FRET efficiency and the diffusion coefficient. Notably, the readout via a change in diffusion can be compared to the theoretical values for the DNA in the bound and unbound states. Furthermore, monitoring the diffusivity serves as an internal quality control check as potential surface-induced artifacts can easily by identified. We further investigated the possibility of performing titrations without the intervention of an operator. The samples were loaded sequentially inside the syringe and were separated using a bubble of air. This approach impeded the use of standard enzymatic oxygen scavenging systems which in turn reduced the overall data quality; nevertheless, the characteristic reduction in FRET efficiency and diffusion coefficient in the samples containing ARF-DBDs was seen. In future experiments, the samples could be loaded in an inert environment (e.g. inert gas atmosphere), to suppress the presence of oxygen between the samples and enable the usage of effective oxygen scavenger systems.

The diffusion coefficients *D* of stochastically labeled AtARF5-DBD and AtARF1-DBD were then used to determine their state of oligomerization in free solution. The values found were compatible with a situation in which the DBDs are present mainly as monomers whereas in crystal structures ARF-DBDs are always present as dimers. This apparent discrepancy serves as a stark reminder that static images of crystal structures, although extremely useful, fail to capture the dynamic nature of biological processes. We further note that the analysis of the displacements along the flow inside the nanochannels allows detecting protein aggregation, if present, which would lead to a slowly diffusing population characterized by advection. As for the individuation of possible surface-induced artifacts, this is a quality control of the sample that is embedded into the methodology.

## Conclusion

In this work we used SMFD inside parallel nanochannels to study the interaction between ARF-DBDs and between ARF-DBDs and their genomic response element. We showed that our approach is able to report on ARF binding by detecting changes in the FRET efficiency similar to experiments with immobilized DNA samples, while obtaining additional information in the form of the displacements distributions. The analysis of this distribution is able to identify the population of absorbed molecules and to separate it from the population of potential aggregates. Moreover, the obtained diffusion coefficient can be used to infer the nature of the tracked molecule/complex. Taken together, our approach allowed us (I) to check for surface induced artifacts and sample quality *during* the experiment, (II) to confirm the results coming from changes in *E*\* with a second readout (*D*) and (III) to infer the oligomerization state of ARF-DBDs in solution. In general, the combination of information from smFRET measurements together with displacement analysis makes SMFD inside nanochannels a powerful method to study protein-protein and protein-DNA association and dissociation.

## Authors’ Contributions

The general research questions were proposed by Dolf Weijers (D.W.) and Johannes Hohlbein (J.H.). J.H. and Mattia Fontana (M.F.) proposed the methodology, defined the experimental design. Simon Lindhoud and Willy van den Berg purified and labelled the proteins. Šarūnė Ivanovaitė and M.F. performed the experiments. M.F. wrote and adapted the software for data analysis and analyzed the data. M.F., J.H. and D.W discussed the content of the manuscript. M.F. wrote the draft manuscript and produced the figures. M.F., J.H. and D.W. edited and finalised the manuscript.

## Data availability

The experimental raw data is currently available upon request and will be made available on https://zenodo.org.

## Acknowledgements

We would like to thank all our colleagues at the Laboratory of Biophysics and the Laboratory of Biochemistry for helpful discussions.

